# Kairos infers *in situ* horizontal gene transfer in longitudinally sampled microbiomes through microdiversity-aware sequence analysis

**DOI:** 10.1101/2023.10.24.563791

**Authors:** Connor L. Brown, Yat Fei Cheung, Haoqiu Song, Delaney Snead, Peter Vikesland, Amy Pruden, Liqing Zhang

## Abstract

Horizontal gene transfer (HGT) occurring within microbiomes is linked to complex environmental and ecological dynamics that are challenging to replicate in controlled settings. Consequently, most extant studies of microbiome HGT are either simplistic experimental settings with tenuous relevance to real microbiomes or correlative studies that assume that HGT potential is a function of the relative abundance of mobile genetic elements (MGEs), the vehicles of HGT. Here we introduce Kairos as a bioinformatic tool deployed in nextflow for detecting HGT events “*in situ,*” i.e., within a microbiome, through analysis of time-series metagenomic sequencing data. The *in-situ* framework proposed here leverages available metagenomic data from a longitudinally sampled microbiome to assess whether the chronological occurrence of potential donors, recipients, and putatively transferred regions could plausibly have arisen due to HGT over a range of defined time periods. The centerpiece of the Kairos workflow is a novel competitive read alignment method that enables discernment of even very similar genomic sequences, such as those produced by MGE-associated recombination. A key advantage of Kairos is its reliance on assemblies rather than metagenome assembled genomes (MAGs), which avoids systematic exclusion of accessory genes associated with the binning process. In an example test-case of real world data, use of assemblies directly produced a 264-fold increase in the number of antibiotic resistance genes included in the analysis of HGT compared to analysis of MAGs with MetaCHIP. Further, *in silico* evaluation of contig taxonomy was performed to assess the accuracy of classification for both chromosomally- and MGE-derived sequences, indicating a high degree of accuracy even for conjugative plasmids up to the level of class or order. Thus, Kairos enables the analysis of very recent HGT events, making it suitable for studying rapid prokaryotic adaptation in environmental systems without disturbing the ornate ecological dynamics associated with microbiomes. Current versions of the Kairos workflow are available here: https://github.com/clb21565/kairos.

## Introduction

Horizontal gene transfer (HGT) facilitates bacterial adaptation in the face of shifting selective pressures. Many clinically-important antibiotic resistance genes (ARGs) have achieved global dissemination through HGT of ARG-bearing mobile genetic elements (MGEs).(R. et al., 2018; U.S. Department of Health and Human Services, 2019; United Nations Environment Programme, 2023) Examining HGT in the context of microbiomes has the potential to yield valuable insights regarding the ecology and evolutionary dynamics of bacterial populations, with especially important implications for antibiotic resistance. For example, HGT of broad host-range MGEs is well documented in the human gut(Brito, 2021; Forster et al., 2022) and has been found to mediate transfer of ARGs across broad phylogenetic ranges, including between gut commensals and potential pathogens.(de Nies et al., 2022; Stecher et al., 2012) Human and animal guts are suspected to be a particularly critical venue for the evolution of resistance in pathogens as clinical concentrations of antibiotics are unlikely to be encountered elsewhere.(Bengtsson-Palme and Larsson, 2016; Gullberg et al., 2011; Larsson and Flach, 2022) However, the environment, and particularly wastewater, is increasingly being recognized for its potential to facilitate the emergence of novel ARGs due to the coalescence of extremely high genetic diversity, MGEs, and selective agents.(Berglund et al., 2023; Ebmeyer et al., 2021)

Increased understanding of the ecological dynamics of HGT in complex environmental microbiomes such as sewage, and the wastewater treatment plants (WWTPs) that treat sewage, could aid surveillance and intervention efforts.(Moralez et al., 2021) For example, the operational parameters of WWTPs are extensively monitored, as required by law. Such monitoring data are essential to adjusting operational conditions as needed and ensuring that performance meets minimum standards of treated water quality prior to discharge. Developing a predictive understanding of bacterial HGT in WWTPs could further enable convenient and synergistic adjustments to operational decisions that could also mitigate unregulated contaminants of concern found in sewage, including antimicrobial resistance determinants. However, no reliable bioinformatic tools exist for monitoring HGT over short timescales in complex microbiomes, such as those represented by WWTPs.(Brito, 2021) Typical approaches include *in vitro* systems with model organisms or analysis of isolate whole genome sequence (WGS) data(Ding et al., 2022; Hutinel et al., 2021; Li et al., 2022), which are unlikely to capture ecological dynamics. Thus, there is a need for tools for tracking HGT that effectively capture the complex interplay between microbial ecology and HGT under real-world conditions.

MetaCHIP(Song et al., 2019), the first such effort towards specifically profiling microbiome-scale HGT, leverages metagenome assembled genomes (MAGs) for HGT detection. While well suited for identifying distant (i.e., older) HGT events, the dependency on MAGs poses several challenges, especially when investigating recent HGT events. It has been shown previously that the accessory genome is particularly difficult to bin accurately when multiple strains of the same species are present.(Maguire et al., 2020; Meziti et al., 2021) This is in part because some portions of the genome are common among strains (core regions) while others (accessory regions) are strain-specific. The result of this is that core and accessory regions display different depth profiles, which makes it challenging to successfully capture both the core and accessory regions in a MAG.(Meziti et al., 2021) Unfortunately, this problem is only exacerbated in the case of mobile ARGs and MGEs, both of which are by definition associated with the accessory genome.(Mazel, 2006; Oliveira et al., 2017)

Here we introduce Kairos as a bioinformatic tool for microbiome-level HGT analysis that addresses many of the above limitations. We further propose a framework of “*in situ* HGT” inference, aiming to provide objective criteria for inferring HGT events occurring within defined windows of time using time series metagenomic sequencing data. The *in situ* framework provides a means to assess whether the chronological occurrence of potential donors, recipients, and putatively transferred regions could plausibly have arisen due to HGT in the sampled period. The centerpiece of Kairos, the Kairos assess workflow, leverages a novel competitive read alignment method that is capable of distinguishing between even very similar genomic sequences. Notably, our methodology is applicable to any longitudinally sampled microbiome for which a reasonable sample of gene contexts can be obtained, thus enabling the potential for retrospective analysis of metagenomic datasets with simplified experimental designs.

## Methods

### Kairos and *in situ* HGTs

Kairos is a nextflow(Di Tommaso et al., 2017) pipeline that integrates multiple tools and python scripts to identify, score, and visualize potential HGTs from a metagenomic assembly. If provided sequencing reads, assemblies, and metadata relaying information about longitudinal aspects of the data, it also can identify potential *in situ* HGT events. We define *in situ* HGT in this context as any putative HGT event for which the chronological occurrence of predicted transferred regions, hosts, and recipients, display patterns of abundance or presence/absence consistent with the event having occurred within the sampled space of the microbiome in question.

Kairos first identifies potential HGTs as identical genes/open reading frames (orfs) shared by the input contigs that have different taxonomic classifications. Subsequent steps assess the bioinformatic support for a given potential HGT and provide the means to assess whether a potential HGT may have occurred *in situ* given a set of longitudinally sampled metagenomes. A complete, step-by-step workflow is described in the supplementary methods and methods below.

### Kairos derep-detect workflow

The Kairos derep-detect workflow takes a set of contigs as input and identifies, scores, and visualizes the potential HGTs **(Fig. 1, Supplementary Methods 1).** The first task in the derep-detect workflow is to identify orfs from a set of contigs. Protein sequences or orfs are predicted using prodigal(Hyatt et al., 2010) (-p meta) and then clustered using mmseqs(Steinegger and Söding, 2017) (coverage of ≥30% and identity of ≥99 %). The orfs predicted from the contigs are also annotated for MGE hallmarks (i.e., mobileOGs) from mobileOG-db(L. et al., 2022) using diamond(Buchfink et al., 2014) (--id 30 --evalue 1e-5) and for ARGs using deepARG-db(Arango-Argoty et al., 2018) (--id 80 --evalue 1e-10 --query-cov 0.6) **(Supplementary Methods 2)**. The user is also able to provide their own database of target genes which will be likewise scored as ARGs are.

**Fig. 1.**
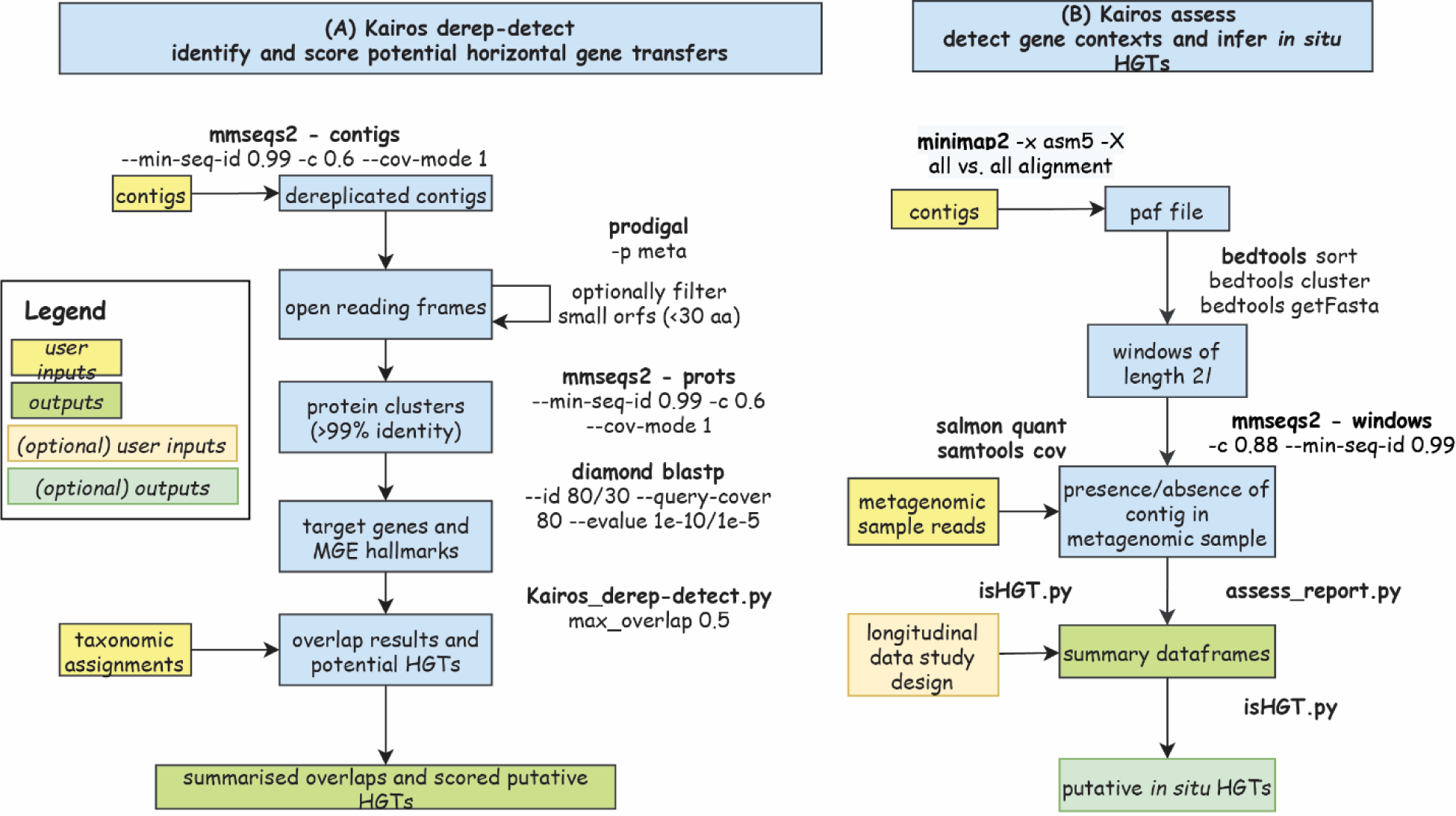
Overview of the Kairos derep-detect and assess workflows for profiling m 94 icrobiome-level HGT via analysis of assembled metagenomic sequences. (A) The Kairos derep-detect workflow takes contigs (capturing a reasonable sample of target gene contexts) and taxonomy assignments as input and produces a list of identical open reading frames (orfs) shared among the contigs and a summary of potential HGTs. (B) The Kairos assess workflow takes contigs and multiple short read samples and produces assessments of contig presence/absences across the set of samples. If provided with additional information regarding the study design, it can infer putative *in situ* HGTs. All settings displayed are default values and are able to be specified by the user.

Optionally, contigs may be dereplicated by calculating the proportion of shared orfs between two contigs. If so, contigs with ≥50% shared orfs relative to the smaller contig (i.e., 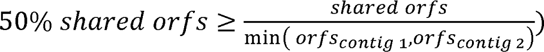 are potential duplicates by default. Clusters are dereplicated by selecting the member with the largest number of orfs as the representative. In the case of ties, one of the tied cluster members are randomly selected. The number of contexts ascribed to a gene is thus the number of dereplicated contigs with the gene.

### Defining potential HGTs

We define any given contig (referring to any contig, scaffold, extracted window from a genome, or other subsection of a genome):

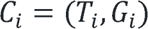

where *T_i_* is the user supplied taxonomic annotation of the contig *C_i_*and *G_i_* is the set of genes on the contig, where *G_i_* = {*g*_1_, *g*_2_,..., *g_n_*}. Two contigs, *C_j_* and *C_k_*, sharing identical genes would be a potential HGT if:

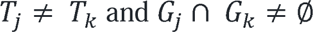

### Scoring potential HGTs

Potential HGTs (as represented above) are ranked according to gene content features **(Supplemental Methods 2)**. First, potential HGTs that involve MGE hallmark genes, such as those aggregated by mobileOG-db, are considered to be more plausible and thus a potential HGT associated with a mobileOG receives a score of 1 and otherwise 0. The mobileOG can either directly be the shared gene or can simply co-occur with the shared gene on one or both of the contigs. In the latter case, the orf matching a mobileOG must be within 5,000 bp of the putatively transferred gene. This distance should be sufficient to be inclusive of co-occurrences with insertion sequence elements, integrative elements, or transposons.(Liu et al., 2019; Ross et al., 2021; Siguier et al., 2015) In addition, a score of 1 is applied if the putatively transferred orf aligns to one of the target database sequences (deepARG-db by default).

### Visualizing potential HGTs

Visualization of putative HGTs is powerful for assessing biological plausibility. The visualize workflow annotates a set of potential HGTs using prokka and visualizes them using clinker(Gilchrist and Chooi, 2021) (**Fig. 2**). The output html files are interactive and can be modified to the user’s preference.

**Fig. 2.**
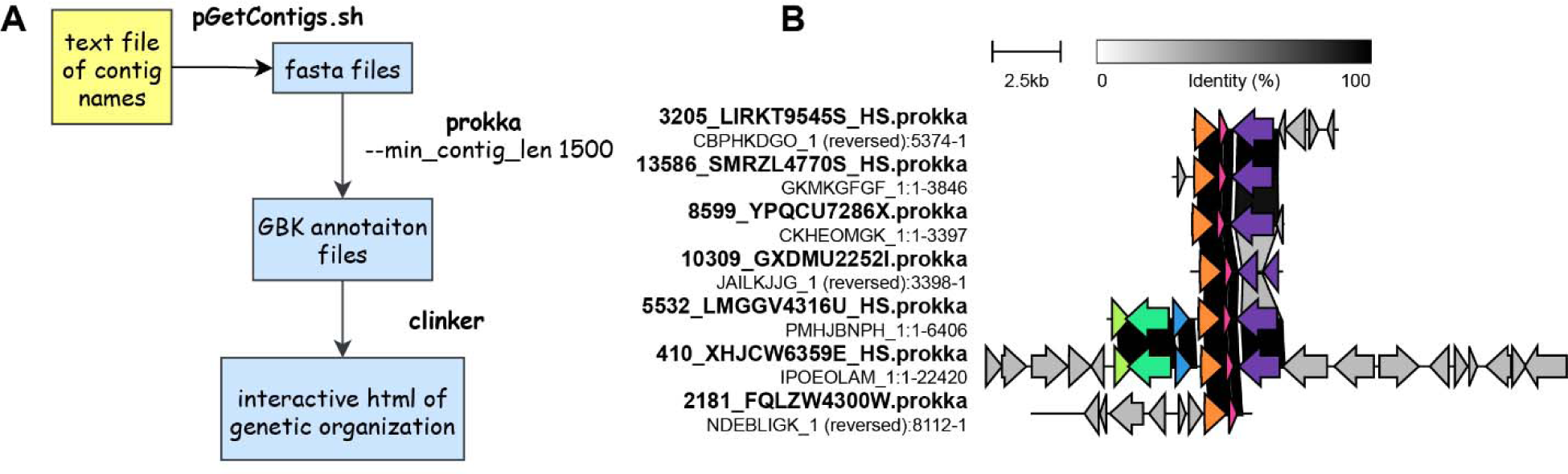
Visualizing potential HGTs provides a powerful means for assessing biological plausibility. (A) Visualization workflow implemented in Kairos as a supplementary script takes in user-supplied text files of contigs to be visualized, extracts them, and produces annotations and visualizations via clinker.(Gilchrist and Chooi, 2021) (B) Example visualizations produced using clinker.

### Inferring in situ HGT events

We define *in situ* HGT as any instance of gene sharing between two contigs with different taxonomic assignments wherein the paired contigs display patterns of presence/absence consistent with an HGT event occurring during a sampled period. Inferring *in situ* HGT events from a longitudinally sampled microbiome is performed using generic and case-specific hypotheses for each instance of potential HGT (**Fig. 3**). For example:

> *H_01_* = The HGT-associated insertion/deletion already existed in the microbiome at a previous time point and thus could not be due to recombination within the period between samplings.

**Fig. 3.**
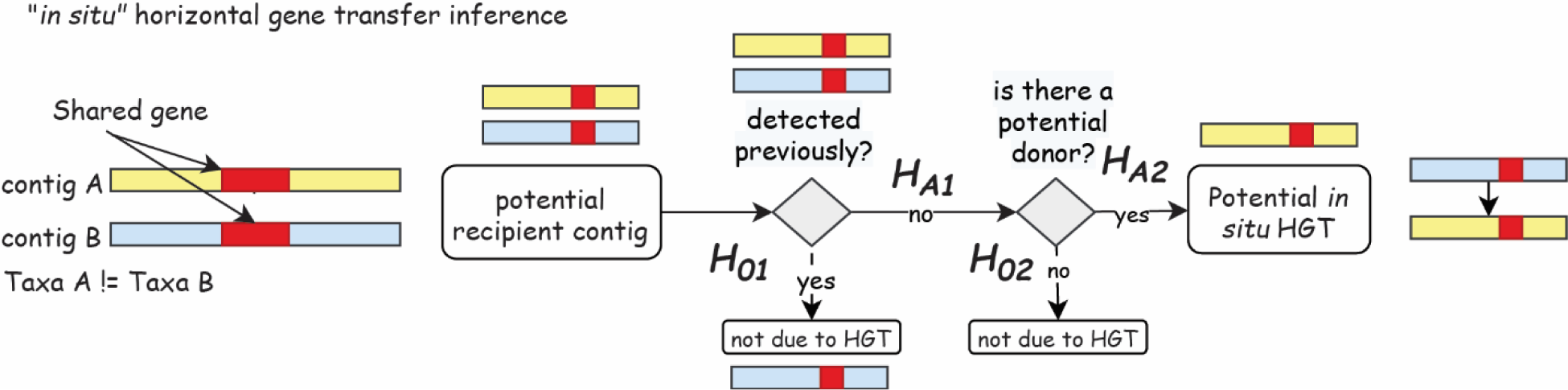
A framework for inferring “*in situ”* HGT events from longitudinal metagenomic data. We propose a framework for inferring HGT occurring within a sampled period of a microbiome (i.e., *in situ*). The potential for *in situ* HGT is assessed by evaluating a set of hypotheses regarding the chronological occurrence of potential donors and recipients in order to determine whether the observed gene co-occurrence could have plausibly arisen within the sampled period.

> *H_A1_ =* The HGT-associated insertion/deletion was not detectable in the microbiome at a previous time point and therefore could have arisen as a result of recombination within the span of the experimental period.

> *H_02_* = Neither contigs comprising the HGT-associated insertion/deletion were detectable at any previous timepoint, implying that there is no detectable donor.

> *H_A2_ =* One of the contigs comprising the HGT-associated insertion/deletion was detectable at a prior timepoint suggesting that the HGT-associated insertion/deletion could have arisen as a result of recombination within the span of the experimental period.

These hypotheses can be amended as additional null hypotheses arise, or as relevant to the experimental design in question.

### Kairos assess workflow provides sensitive detection of contigs associated with potential HGT

The Kairos assess workflow identifies and then uses boundary regions defined by the edges of an alignment between two contigs **(Fig. 1, Supplementary Methods 3, Fig. S1)** to further investigate the potential HGTs. All vs. all alignment of a set of contigs with potential HGTs is performed using minimap2(Li, 2018) (-x asm5 -X). Edge regions, defined by coordinates of (alignment-start ± length *l* and alignment-end ± length *l*, default of 75 bps) are written to bedfiles that are then sorted, clustered, and extracted from the contigs using bedtools.(Quinlan, 2014) Edges are dereplicated using mmseqs (identity ≥99% and coverage = 88%). Short reads are mapped to the dereplicated edges using salmon(Patro et al., 2017) quant and the presence/absence of each region of variation are assessed by counting the number of reads mapping to each boundary region passing quality filtering (100 bp minimum alignment length, i.e., samtools view -m 100). By default, a minimum of one read is taken as evidence of the locus being present. The minimum alignment setting of 100 bp ensures that at least 25 bp of the unique portion of the locus is present (hence, it is a ‘competitive’ reads mapping approach). Results of the read mapping are summarized using samtools(Danecek et al., 2021) coverage (using default parameters). Read mapping results are extended to apply to edge cluster members by combining the output of samtools coverage with the edge cluster table. The presence/absence of structural variations are determined by counting the proportion of distinguishing boundaries detected to total distinguishing boundaries in the contig (≥90% of distinguishing boundaries must be detected).

### Longitudinally sampled sequencing batch reactors

Sequencing batch reactors (SBRs) were operated using influent recovered from a local municipal WWTP and large urban hospital in Illinois. Extended details of SBR operation can be found elsewhere.(Brown et al., 2023; Maile-Moskowitz, Ayella,Connor Brown, Latania Logan, Kang Xia, Amy Pruden, 2023) The SBRs were seeded with activated sludge from the corresponding municipal WWTP and were maintained for a period of weeks prior to reaching steady-state operation (i.e., stable removal of organic carbon). For the following three weeks, samples were collected for culture of antibiotic resistant pathogens (*Klebsiella pneumoniae, Escherichia coli,* and carbapenemase producing Enterobacterales (CPE). This produced a catalogue of 456 isolates in addition to 111 Illumina shotgun metagenomes of influent, effluent, and AS, and 36 nanopore long read samples. AS and influent samples were sequenced to approximately 5 Gbp per sample and effluent to 3 Gbp per sample. A subset of AS and influent samples (*n* = 6) were also subjected to deep sequencing (mean 36 Gbp per sample).

### Assembly of an MGE and resistance gene catalogue

We contrasted MAG-based inferences with a catalogue of contigs with a catalogue of MGEs and resistance genes created in a parallel study.(Brown et al., 2023) Briefly, multiple hybrid assembly strategies were performed using short Illumina reads and long minION nanopore reads to improve recovery of informative resistance gene contexts. Briefly, individual samples were assembled using OPERA-MS(Bertrand et al., 2019) (--contig-len-thr 1000 –long-read-mapper minimap2) and hybridSPAdes(Antipov et al., 2016) (metaspades.py with default settings). OPERA-MS was used for all coassemblies, including individual reactors (e.g., 10%-1) across all timepoints, coassembly of all ML samples, and of samples partitioned by treatment (i.e., ± hospital effluent or 10% vs. 0%). All assemblies/coassemblies were searched for RGs and MGE hallmark genes. Protein sequences were predicted using prodigal (-meta) and queried against experimental sequences in BacMet v2,(Pal et al., 2014) CARD v3.0.7,(Alcock et al., 2020) and mobileOG-db beatrix-v1.6(L. et al., 2022) using diamond(Buchfink et al., 2014) blastp (-id 90% -e 1e-10). For subsequent contextual analysis, only those contigs with a hit from one of the databases was retained.

### MAG recovery and dereplication

Assemblies produced in the creation of the MGE and resistance gene catalogue were further binned using both MetaBat2(Kang et al., 2019) and MaxBin.(Wu et al., 2016) Creation of sorted bam files was performed using minimap2(Li, 2018) read alignment (-x sr) of the corresponding short read samples. Only the coassembly of all samples (excluding deeply-sequenced ones) were used for binning using both bbmap, minimap2 and subsequently MetaBat2 and MaxBin. The resulting draft MAG collection was dereplicated using dRep v. 2 with default settings.

### *In silico* validation of Kairos assess

We assessed Kairos’s ability to distinguish samples with and without simulated plasmid sequences bearing small differences in sequence (**Supplementary Methods 4**). Sequenced plasmid assemblies were extracted from the assembled WGS of *Aeromonas rivipollensis* ArCPE-VT-1 and *Escherichia coli* EcrMDR-VT-1. We additionally identified two plasmids with >99% ANI from plsdb(Schmartz et al., 2022) using blastn v.2.12.0+ **(Table S1)**. To simulate an insertion, ISEscan(Xie and Tang, 2017) was used to identify copy of IS91, a cut-and-paste type transposable element, from one metagenomic assembly **(Fig. S2)**. The extracted copy of IS91(Berger and Haas, 2001) was inserted into a random position in the WGS-derived and plsdb-derived plasmid sequences **(Fig. S3)**. Simulated chimeric sequences were generated by randomly merging 2,500 bp windows extracted from the plasmid sequences. Strain-level chimeras were those where the source plasmids had >99% ANI (i.e., were derived from a WGS sequence and its closest match from plsdb). More distant chimeras were generated by splicing either WGS with WGS plasmid sequences, or with plsdb with plsdb sequences. Reads were then simulated using in silico seq(Gourlé et al., 2019) (iss generate --seed 1 --cpus 32 --genomes merged_simulated.fasta --abundance uniform --n_reads 1000000 --model NovaSeq --mode kde - -o is_reads) and were spiked into the appropriate test samples **(Supplementary Methods 4, Table S2)** at 1×, 5×, or 10× coverage.

### Evaluating taxonomic classification of bacterial chromosomes, plasmids, phages, and mobile genetic elements

To provide guidance on the conditions that provide reliable taxonomic inferences for contigs, we evaluated taxonomic classification using three different methods (kraken2(Wood et al., 2019) with gtdb,(Parks et al., 2022) kraken2 with the standard reference database (downloaded August 2022), and mmseqs2 taxonomy(Mirdita et al., 2021) using gtdb (v202). We selected a set of 2,178 environment-associated bacteria and archaea from GenBank **(Table S3**) from which we simulated contigs of 500 bp, 1,500 bp, 3,000 bp, and 5,000 bp in size by fragmenting the genomes using seqkit(Shen et al., 2016) and subjected them to taxonomic annotation. In addition, we also assessed the fidelity of taxonomic assignments of MGEs applied to plasmids (COMPASS),(Douarre et al., 2020) integrative elements (ICEberg 2.0),(Liu et al., 2019) and phages (pVOG)(Grazziotin et al., 2017) using only 3,000 bp length fragments and mmseqs2 with gtdb **(Table S3)**. MGEs with genus labels of *Raoutella, Shigella, Mycolicobacterium* were relabeled as *Klebsiella, Escherichia,* and *Mycobacterium*, respectively, consistent with gtdb.

## Results and Discussion

### Kairos enables capture of HGT in the unbinned accessory genome via direct analysis of assemblies

The Kairos derep-detect workflow predicts potential HGTs directly from contigs rather than relying on MAGs. This is in contrast to MetaCHIP, which leverages MAGs for its inferences. Here and throughout, we employ data generated from a controlled and replicated experiment using SBRs, a lab-scale bioreactor commonly employed for replicable simulation of activated sludge wastewater treatment.(Brown et al., 2023; Maile-Moskowitz, Ayella,Connor Brown, Latania Logan, Kang Xia, Amy Pruden, 2023) Sampling of the SBRs took place over three weeks, during which time isolates of multidrug resistant bacteria were collected in addition to samples for shotgun metagenomics using both Illumina and nanopore sequencing platforms.

Comparison of the MAGs, assemblies, and WGS data highlights the strengths and weakness of three options for characterizing microbiome-level HGT (**Fig. 4**). After binning and dereplication(Brown et al., 2023) only 66 ARGs (54 unique) and 3,810 mobileOGs (3,182 unique) were detected in the 876 MAGs vs. 17,954 ARGs (537 unique) and 1,408,559 mobileOGs (91,710 unique) in the MGE/resistance gene catalogue. Thus, use of metagenomic assemblies directly, rather than MAGs, averted a 265-fold loss of resistance gene information.

**Fig. 4.**
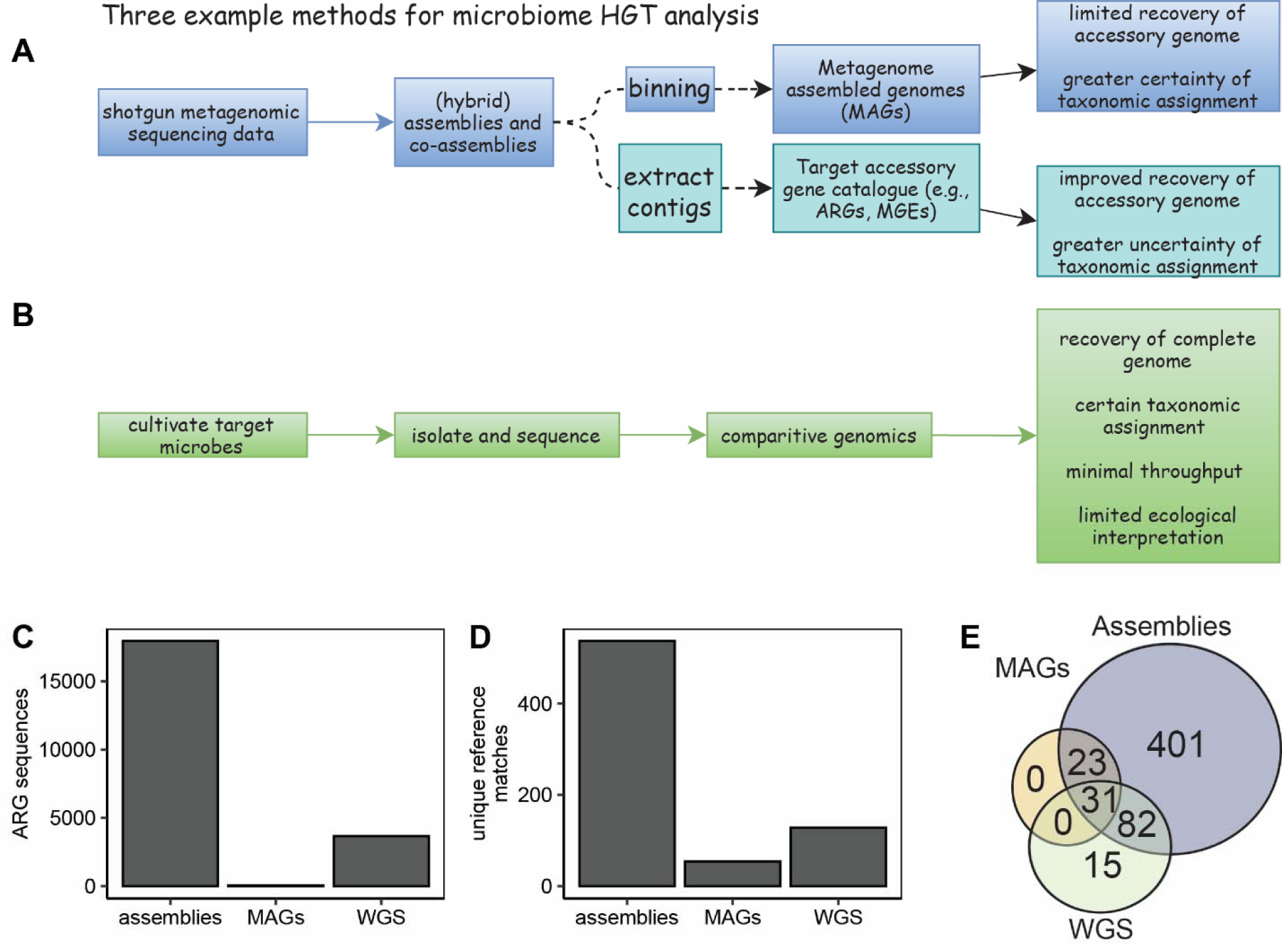
Overview of three different methods for tackling microbiome HGT. (A) Two potential routes to identifying HGT in a microbiome start with assembly of shotgun metagenomic sequencing data and lead to either analysis of binned assemblies, i.e., metagenome assembled genomes (MAGs) or via direct analysis of assemblies. MAGs have taxonomic assignments that are more certain. However, the binning process tends to exclude important accessory genes. Analysis of assemblies directly (i.e., no binning) improves recovery of accessory genes but means less certainty in taxonomic assignments. (B) One alternative approach might be to cultivate and isolate relevant species, for example, drug resistant pathogens, and subject to whole genome sequencing (WGS). While providing certain taxonomic assignments and robust coverage of accessory genes, there is limited throughput, and the process excludes non-culturable organisms. The methods presented here do not comprise an exhaustive list of experimental approaches.(Brito, 2021) (C-E): report results from a lab scale study of activated sludge for which culture and metagenomic data were obtained. (C) A total of 17,954 ARG-encoding orfs were detected in the assembled contigs vs. 66 in MAGs. (D) A total of 573 unique ARG reference sequences were detected in the assembled contigs vs. 54 in MAGs. (E) Assemblies, MAGs, and WGS contain partially overlapping sets of the resistome, with assemblies capturing the most.

Among the 66 ARGs detected in the MAGs, about half (28, 42%) were detected in a MAG with a strain-level taxonomic assignment of *E. coli* D (bin86). This MAG was likely derived from the same clonal lineage as one of the isolates with an ANI value >99.99%. Encouragingly, the MAG-associated ARGs entirely overlapped with ARGs encountered in the WGS of the *E. coli* isolate. However, the MAG lacked 58 ARGs that were associated with the WGS. Further scrutiny reaffirmed that many of the MAG-encoded ARGs were those typically encoded on chromosomes (e.g., genes encoding an AmpC-type beta-lactamase and a TolC outer membrane protein) **(Table S4)**, and thus were unlikely to be constituents of the accessory genome. Notably, 15 ARGs detected in WGS were not present in the metagenome assemblies.

### Concordance of network properties of predicted HGTs across source genomic catalogue and method

We next conducted a parallel comparison of MetaCHIP versus Kairos derep-detect using MAGs and assemblies, respectively. It was noted that when running MetaCHIP, the overall computation time for the bins (876 MAGs totaling 3.19 Gbp with an N_50_ of 16,765) was a little over 2 days on an institutional high performance computing cluster (128 cores with 200 GB memory). The majority of this time was devoted towards the all vs. all blastn step. By contrast, the Kairos derep-detect workflow required about 1 hour. It should be noted that recent versions of MetaCHIP have pivoted from using blastn and substituted it for minimap2.

Overall, the predicted gene sharing networks produced by the two pipelines were similar across tools and target catalogues (i.e., MAGs or contigs) (**Fig. 5A-C**). Over the full range of conditions examined (i.e., Kairos derep-detect workflow applied to the assemblies or MAGs; and MetaCHIP applied to the MAGs), network topology was found to be similar in terms of degree (i.e., the number of edges corresponding to a particular node) and neighborhood connectivity (the average number of edges corresponding to the first order-neighbors) (**Fig. 5D,E**). Notably, Kairos and MetaCHIP agreed in terms of overall rates of HGT (**Fig. 6A**) estimated across different taxonomic strata. However, it was also noted that estimated genus level HGTs were highest when using Kairos with the MAG catalogue. Closer examination of the gene families putatively enriched in HGT predictions produced by Kairos using MAGs revealed the presence of several conserved protein families (e.g., PFAM Sigma70_r2 associated with bacterial RNA polymerase), suggesting that such families may be prone to erroneous classification when using Kairos/MAGs (**Fig. 6B**). By contrast, MetaCHIP likely correctly eliminates them through phylogenetic analysis, which compares single copy gene evolution to putative HGT genes to differentiate HGT from vertical inheritance. While the potential for misclassification of highly conserved protein families by Kairos when MAGs are used as the input data is a duly-noted limitation, it likely could be subverted by excluding HGTs without co-occurring MGE hallmark genes.

**Fig. 5.**
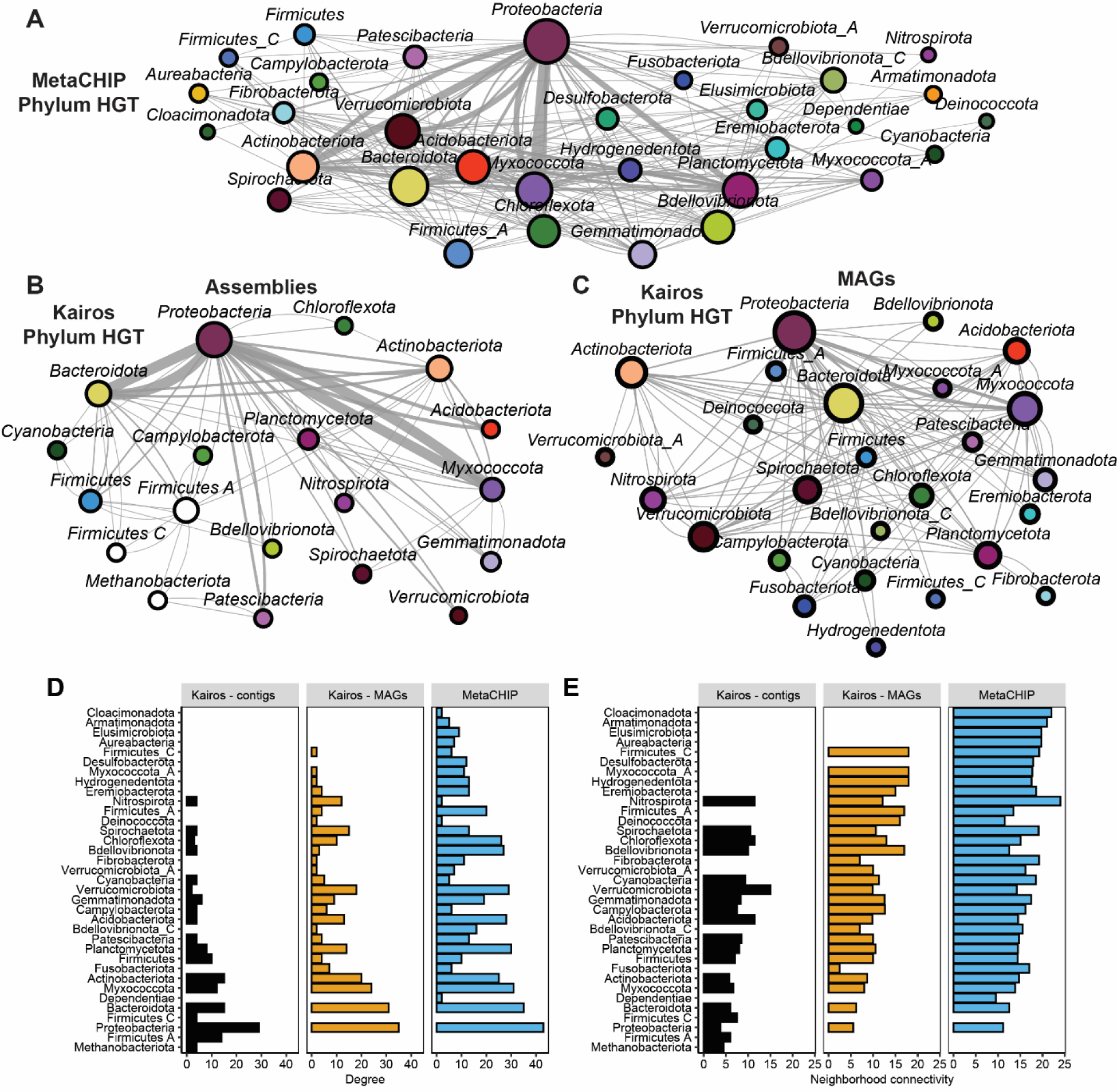
Concordance of network properties of predicted HGTs across source genomic catalogue and HGT prediction approach. Network of MetaCHIP-predicted HGTs weighted by frequency of predicted gene sharing highlights centrality of *Proteobacteria, Verrumicrobiota,* and *Bacteroidia.* (B) Network of Kairos derep-detect predicted HGTs weighted by frequency of predicted gene sharing using the assemblies with taxonomic assignments derived from contigs. (C) Network of Kairos derep-detect predicted HGTs weighted by frequency of predicted gene sharing using the MAGs with taxonomic assignments derived from MAGs. (D) Phylum-level degrees (the number of edges corresponding to a particular node) from networks A-C highlight similarity in topology between the three networks. (E) Phylum-level neighborhood connectivity values (the average number of edges corresponding to the first order-neighbors) again highlight similarities between the three networks. A detailed display of the experimental design is provided **(Fig. S4).**

**Fig. 6.**
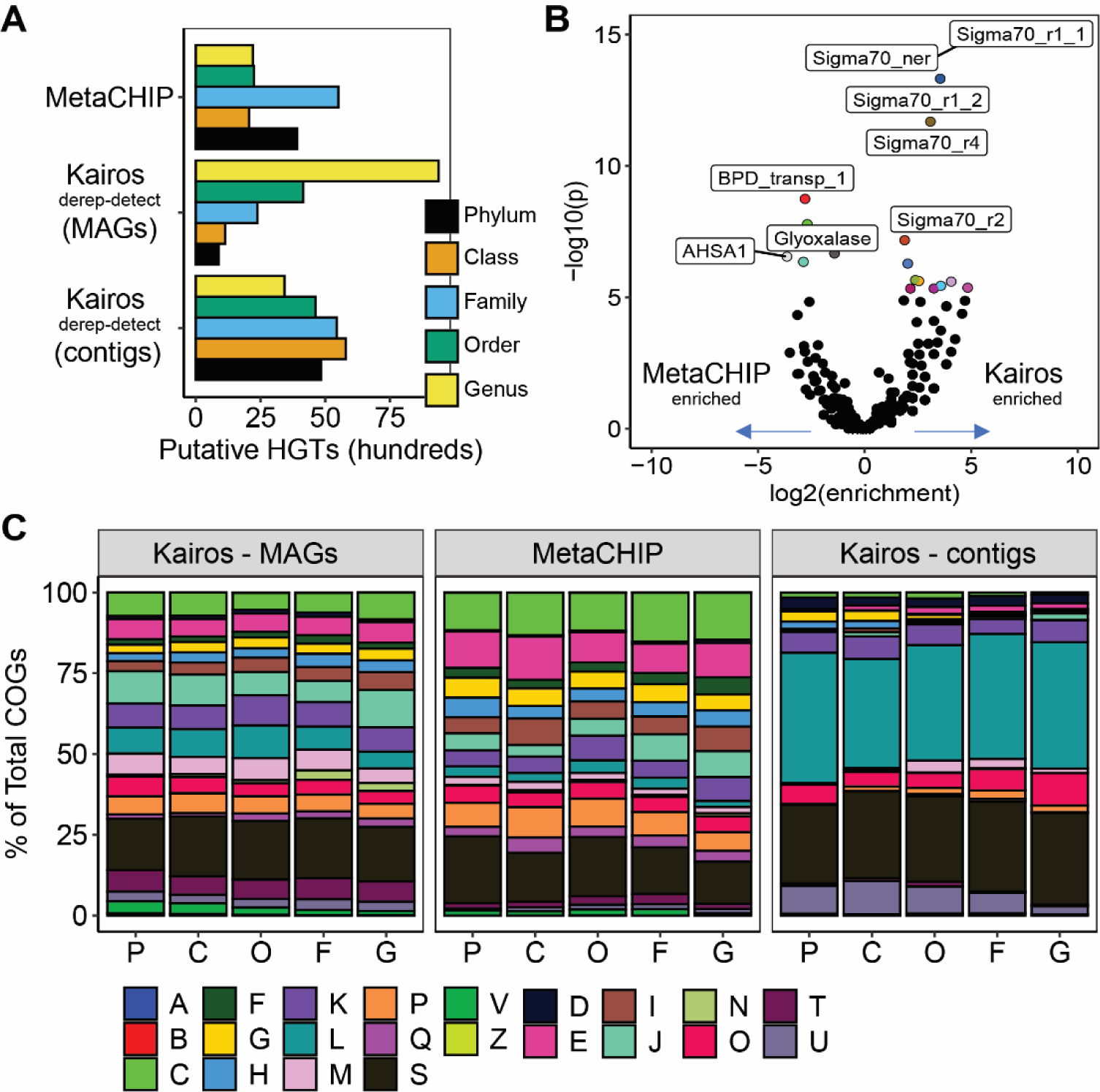
Head-to-head comparison of MetaCHIP and Kairos applied to MAGs and assemblies. (A) Overall number of predicted HGTs partitioned by taxonomic level. Kairos derep-detect when applied to MAGs suggested a high number of HGTs occurring at the level of genus. B) Volcano plot of PFAMs enrichment in either Kairos MAG genus-level HGT predictions (log2(enrichment)>0) or MetaCHIP predictions (log2(enrichment)<0). (C) Comparison of COG categories predicted to be transferred between different methods. X-axis refers to taxonomic level (P: phylum; C: class; O: order; F: family; G: genus).

Examining the functional categories predicted to be transferred by the different methods, the use of assembled contigs had an increased proportion of COG category L (replication, recombination, repair), S (unknown function) and U (secretion/intracellular trafficking) proteins relative to the other two approaches (**Fig. 6C**). By contrast, use of MAGs as input produced more frequent predictions of COG categories C, E, and P, which are linked to energy production, amino acid metabolism, and inorganic phosphate metabolism, respectively. This is likely due to the differences in the genome “fractions” (i.e., core vs. accessory genes) represented by the two catalogues.

### Fidelity of contig taxonomic assignments

An important concern regarding the use of contigs rather than MAGs for HGT inference is to what extent the contig taxonomic annotation is trustworthy. First, predictions of contig taxonomy might be inaccurate due to contamination of the underlying genome database,(Abraham et al., 2023) for example. This could lead to false positive prediction of HGT between genomes of the true and erroneous taxonomic assignments. Second, lack of phylogenetic signal in a contig might result in a low resolution assignment, essentially masking HGT at higher taxonomic levels (e.g., genus or species). Of particular concern are the taxonomic annotations of MGEs which, by definition, have transient associations with individual bacteria. To interrogate what impact these challenges have on yielding accurate taxonomic inferences, we performed a series of experiments examining the efficacy of taxonomic inference using mmseqs2 or kraken2 on simulated contigs derived from chromosomes of environmental bacteria, plasmids, phages, and integrative elements **(Figs S5-7).** Briefly, it was found that taxonomic inferences using mmseqs2 with gtdb as the underlying taxonomic reference database yielded the best performance (i.e., greatest accuracy) for chromosomal sequences (with phylum-level accuracies ranging from 98.72%-99.24%) **(Fig. S5)**. Taxonomic annotation of contigs simulated from MGEs using mmseqs2 displayed a wider range of accuracies **(Fig. S6A)** and higher rates of unclassified sequences **(Fig. S6B).** We additionally observed that contigs derived from conjugative or mobilizable plasmids were less frequently classified (median 90% for non-mobilizable plasmids vs. 80% for conjugative and 56% for mobilizable plasmids at the order taxonomic level). However, conjugative plasmid contigs that were classified had median accuracies above 75% at the level of genus and >90% at the family level **(Fig. S7).** This suggests that if annotation of contigs successfully produces a taxonomic classification, then the taxonomic assignment is generally accurate, even for plasmids. However, it is not possible to know with certainty the host of a plasmid sequence in a metagenome without additional lines of evidence

### Kairos provides sensitive detection of structural microdiversity while reducing the inclusion of chimeric assemblies in the analysis

Analysis of very recent HGT involving recombination of some sort requires consideration of structural microdiversity (i.e., variation in a genomic region of 1 kbp or more) to successfully distinguish between closely related genome sequences with and without a putative recombination event. However, this is difficult to distinguish from chimeric assemblies, i.e., assembled sequences that are derived from more than one genome. Chimeric assemblies are especially problematic in the context of HGT and gene sharing analyses as they can produce false-positive associations between taxa, MGEs, and cargo genes. The Kairos assess workflow addresses this through microdiversity aware sequence analysis (**Fig. 7A-D**). We assessed the competitive read alignment method for its ability to distinguish the presence or absence of plasmid sequences with or without an *in silico* inserted copy of insertion sequence IS91 that was extracted from the assemblies **(Figs. 7E-G, S2-S4)**. The plasmids in question were derived from isolates recovered from the SBRs and close matches to the plasmids in plsdb, for a total of eight plasmid sequences (**Table S1**). In addition, we compared this method to a static breadth of coverage (BoC) cut-off 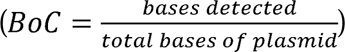 based detection.

**Fig. 7.**
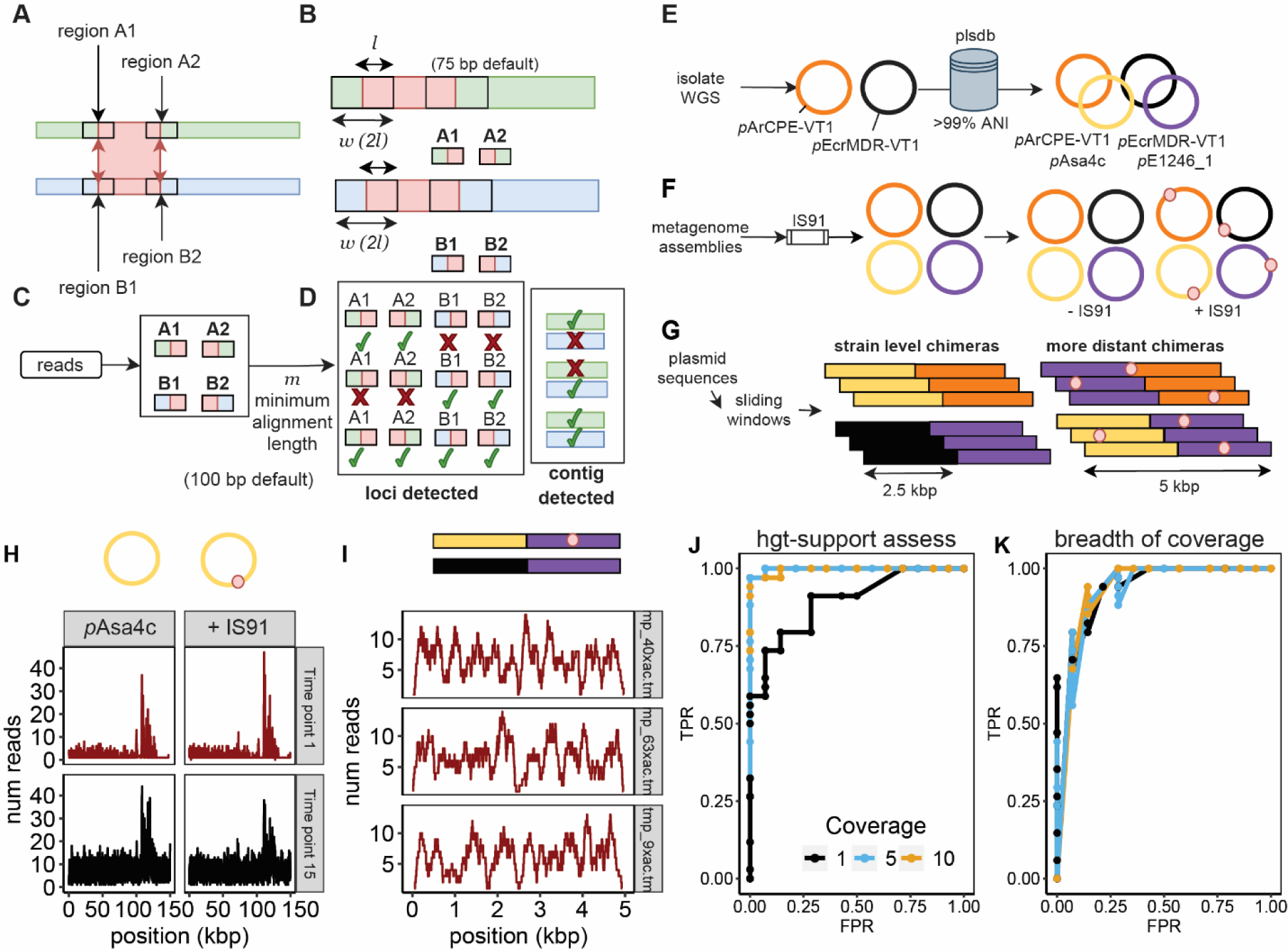
The Kairos assess method provides sensitive detection of mobile element-associated microdiversity through competitive read mapping. (A-D): The Kairos assess method. (A) Two contigs sharing an aligned region with A1/A2 and B1/B2 representing boundary regions of the alignment are identified. (B) Windows of length 2*l* (75 bp by default) in 5’ and 3’ directions on both contigs are extracted. (C) Reads are aligned to the extracted window regions and must meet a minimum alignment length (100 bp by default to ensure a minimum of 25 bp of the unique region is kept). (D) Pattern of distinguishing boundary regions presence/absence (i.e., A1, A2, B1, or B2) is used to infer the presence of the contig they are derived from. Each row represents the different possibilities for two contigs A and B. For example, in the top row, only windows from A are detected and thus contig A is determined to be present (as displayed in the second column). (E) Identification of plasmid sequences (both WGS-derived and public database-derived) for evaluations. (F) Generation of *in silico* insertions using a copy of IS91. (G) Simulation of chimeric fragments. (H) Breadth of coverage is unable to distinguish plasmid strains with or without a copy of IS91. Red color indicates the sample that is mapped did not receive the spike-in. (I) Example coverage profiles of three randomly selected chimeras are indistinguishable from correctly assembled fragment depth profiles (e.g., panel H). (J) ROC curve highlighting the influence on target coverage (1×, 5×, and 10×) on the efficacy of Kairos assess for determining presence/absence of plasmid genomes (created in panels E and F). (K) Breadth of coverage shows worse performance in distinguishing the presence/absence of plasmids with or without IS91.

As expected, BoC was a poor indicator of plasmid presence or absence as all sequences retained ≥ 50% BoC across all samples used for these analyses **(Fig. 7H,I, Table S5)**. By contrast, the support method provided near perfect detection of the plasmids (precision = 1 and recall = 0.97) at 10× coverage while 5× coverage displayed slightly superior performance (**Fig. 7J**). However, even at lower coverages, Kairos assess maintained a greater accuracy than did BoC-based classification (**Fig. 7J,K**). We hypothesized that the support method would eliminate chimeric assemblies from analysis because chimera-derived loci would be unlikely to yield a sufficient number of ≥100 bp alignments. To test this, we combined 2.5 kbp fragments of the isolate-derived plasmids with 2.5 kbp fragments of plasmids derived from plsdb, simulating a strain-level chimeric assembly. We also combined 2.5 kbp fragments of the *E. coli* and *A. rivipollensis* plasmids containing the IS91 copy simulating chimeric assembly of more distantly related plasmids, in part to mirror chimeric assemblies due to shared copies of MGEs. This experiment was conducted using 10× coverage to maximize the potential for false positive detection of chimeric fragments. This produced encouraging results, with 94.67% accuracy for strain-level chimeras (e.g., *A. rivipollensis* + *A. rivipollensis*) and 97.60% accuracy for chimeras constructed from divergent plasmids (e.g., *A. rivipollensis* + *E. coli*). Despite not completely eliminating chimeric fragments entirely, the method demonstrated an overall tendency of exclusion. This suggests that Kairos assess provides sensitive detection of contigs representing structural microdiversity, while simultaneously diminishing chimeric assemblies.

### *In situ* HGT analysis for incorporating ecology into environmental HGT models and hypothesis generation

The framework proposed was specifically configured to enable HGT-relevant hypothesis testing using longitudinally sampled microbiomes. An initial application of the *in situ* HGT framework revealed multiple putative pathways and ecological dynamics of ARG transfer in activated sludge linked to fluctuations in antibiotic levels(Brown et al., 2023). However, Kairos is unlikely to detect instances of conjugation that did not also involve some form of recombination. Conjugation is typically mediated through physical interactions between cells through the activity of a protein supramolecular complex that translocates single-stranded DNA across donor and recipient membranes.(Costa et al., 2021) This generally results in replicative transfer of an identical copy of the MGE into the recipient cell,(Humbert et al., 2019) which would not be distinguishable on the basis of gene content. On the other hand, Kairos is especially suited to address HGT associated with recombination, such as that posed by transposable elements and cargo elements of conjugative MGEs, and transduction. Indeed, initial applications of Kairos recently suggested the transduction of macrolide resistance gene *mphA* across classes *Myxococcia* and *Polyangia*, two species of the phylum *Myxococcota.* The gene itself appears to have originated from a *Proteobacteria* of the order *Xanthomonadales.* Thus, Kairos is able to address modes of HGT beyond conjugation, a functionality that has been critically lacking in existing approaches.(Brito, 2021; R. et al., 2018)

Including additional DNA sequencing data types in the analysis, such as complimentary long read or Hi-C sequencing, could help to further improve detection microbiome-level HGT, but this is not a requirement for Kairos. One note of caution is that metagenomic assembly is notoriously prone to error due to the inherent complexity of environmental microbiomes, which challenges computational algorithms. We previously assessed multiple means of short-, long-, and hybrid-assembly and found that hybrid assembly greatly improved the accuracy and length of metagenomic assemblies associated with wastewater, a complex environmental microbiome. (Brown et al., 2021) Our results also suggested that contigs with greater coverage (>5× coverage) were less likely to be chimeras, although displayed increased rates of insertions and deletions. In the future, we envision that improved methods for assembly graph mining could enable more exhaustive production of assembled genomic catalogues directly from short read metagenomes.

Comprehensively identifying and quantifying key microbial ecological factors driving microbiome-level HGT remains a critical frontier towards characterizing microbial evolution across a suite of different domains. While, like any other method, metagenomic sequencing has inherent limitations, the *in situ* framework presented here achieves its intended purpose of generating hypotheses to support the development of models that characterize potential HGT pathways at the microbiome-scale. The represents a substantial step forward towards understanding such complex phenomena *in situ*, relative to extrapolating from simplistic experiments.

