## Supplementary Information for "Kairos infers *in situ* horizontal gene transfer in longitudinally sampled microbiomes through microdiversity-aware sequence analysis"

### Supplementary Methods 1: Kairos derep-detect: orf-based dereplication and overlap detection.

Scripts: https://github.com/clb21565/kairos /blob/main/scripts/descriptions.MD

***Default settings are listed in parentheses***

1. Predict open reading frames (orfs) using prodigal (1) (-p meta)
2. Cluster orfs using mmseqs (2) (coverage of ≥60% and identity of ≥99 %)
3. Cluster contigs by proportion of shared orfs (minimum of 50% by default)
   a. Contigs with ≥50% shared orfs relative to the smaller contig (i.e., $50\% shared orfs\geq\frac{shared orfs}{\min\left( {orfs}_{contig 1},{orfs}_{contig 2} \right)})$ are potential duplicates.
   b. Clusters are dereplicated and the member with the largest number of orfs is selected as the representative.
   c. In the case of ties, one of the tied cluster members are randomly selected.
   d. The number of contexts ascribed to a gene is thus the number of dereplicated contigs with the gene.

### Supplementary Methods 2: open reading frame annotation, and potential HGT scoring

1. User supplied taxonomic assignments are taken as a TSV with column IDs of “contig” and “lineage”
   a. We recommend using mmseqs taxonomy(3), as in the command mmseqs: taxonomy contigs $GTDB assignments tmpFolder --tax-lineage 1 --majority 0.5 --vote-mode 1 --lca-mode 3 --orf-filter 1)
2. orfs produced from user supplied contigs are aligned to mobileOG-db(1) (v. 1.6.1) and deepARG-db (v2.0)(4) using diamond blastp (mobileOG-db: --pident 30 --evalue 1e-5; deepARG-db: --pident 80 --evalue 1e-10 --query-cover 80). The user may also specify a supplementary database for analysis.
3. Taxonomy information, protein annotation information, and overlap detection from previous steps are integrated and potential HGTs are scored. Summary data frames are written.

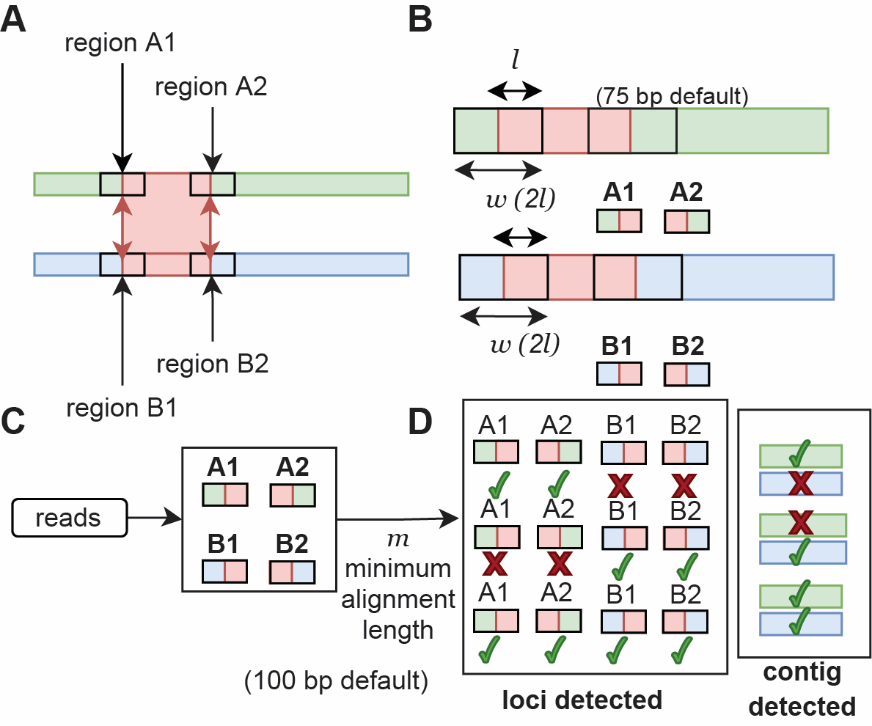

#### Fig. S1. Kairos competitive read mapping step in the Kairos assess workflow. (A) Two contigs sharing an aligned region with A1/A2 and B1/B2 representing boundary regions of the alignment are identified. (B) Windows of length 2*l* (75 bp by default) in 5’ and 3’ directions on both contigs are extracted. (C) Reads are aligned to the fragments with a minimum alignment length (100 bp by default). (D) Pattern of distinguishing loci presence/absence (i.e., A1, A2, B1, or B2) is used to infer the presence of the contig they are derived from.

### Supplementary Methods 3: Kairos assess workflow

1. Deduplicated contigs are aligned to one another using minimap2 (-x asm5 -X 5000)
2. Edge regions, defined by coordinates of (alignment-start ± length *l* and alignment-end ± length *l*, default of 75 bp) are written to bedfiles that are then sorted, clustered, and extracted from the contigs using bedtools(5)
3. Edges are dereplicated using mmseqs (identity ≥99% and coverage = 88%)
4. Short reads are mapped to the dereplicated edges using salmon quant and the presence/absence of each region of variation are assessed by counting the number of reads mapping to each locus passing quality filtering (100 bp minimum alignment length, i.e., samtools view -m 100)
   a. By default, a minimum of one read is taken as evidence of the locus being present. The minimum alignment setting of 100 bp ensures that at least 25 bp of the unique portion of the locus is present.
5. Results of the reads mapping are summarized using samtools coverage(6) (using default parameters).
6. Reads mapping results are extended to apply also to edge cluster members by combining the output of samtools coverage with the edge cluster table.
7. The presence/absence of structural variations are determined by counting the proportion of distinguishing loci detected to total distinguishing loci in the contig (≥90% of distinguishing loci must be detected).

### Supplementary Methods 4: Evaluating Kairos assess

To assess the sensitivity of the Kairos assess workflow, *in silico* plasmid variant sequences were generated. Two plasmids were extracted from the assemblies of *Escherichia coli* strains EcrMDR-VT-1 and *Aeromonas rivipollensis* strain ArCPE-VT-1 isolated from the reactors (**Table S1**). Close relatives of these plasmids in public databases were detected by mapping the sequences against plsdb (7) using blastn and the top-scoring match was extracted for both. A single IS91 insertion sequence element was predicted from a small set of randomly selected contigs from the reactor assemblies using ISEscan and inserted into each of the isolate-derived and public database-derived sequences at random points resulting in eight total plasmid sequences (four with random insertions and four without). For the *E. coli* plasmid, the closest reference plasmid sequence was NZ_CP025574.1 (with 99.5789% average nucleotide identity) while the closest match to the *A. rivipollensis* plasmid was NZ_CP054295.1 (with 99.83% average nucleotide identity), corresponding to NZ_KT033470.1 *Aeromonas salmonicida* subsp. *salmonicida* strain JF2267 plasmid pAsa4c.

#### Table S1. Plasmids used in the present study for *in silico* validation.

| **Table S1. Plasmids used in the present study for *in silico* validation.** | | | |
| --- | --- | --- | --- |
| Sequence name | With IS91 | Strain | Reference |
| AR-real | no | Aeromonas rivipollensis ArCPE-VT-1 | This study |
| AR-real | yes |  |  |
| EC-plsdb | no | NZ_CP025574.1 Escherichia coli strain E-1246 plasmid pE1246_1 | https://journals.asm.org/doi/10.1128/mBio.00347-19 |
| EC-plsdb | yes |  |  |
| EC-real | no | Escherichia coli EcrMDR-VT-1 | This study |
| EC-real | yes |  |  |
| AR-plsdb | no | NZ_KT033470.1 Aeromonas salmonicida subsp. salmonicida strain JF2267 plasmid pAsa4c | https://pubmed.ncbi.nlm.nih.gov/27812409/ |
| AR-plsdb | yes |  |  |

>615_LNBYT5954Z_HS_10644_12836_+ IS91_50

GGTAATCGCCCTGCGCGCCGTGGAAACCATCGACTTCATGACCGCGCACTGGGCGCACCT

GCCGTATGAGTTTCTGGGCAGGGTCTCCAACCGCATCATTAACGAACTGCGCGGCGTCTC

GCGCGTGGTCTACGACATCTCCGGCAAGCCACCGGCGACGATTGAGTGGGAGTGACGGGC

ACTGGCTGGCAATGTCTAGCAACGGCAGGCATTTCGGCTGAGGGTAAAAGAACTTTCCGC

TAAGCGATAGACTGTATGTAAACACAGTATTGCAAGGACGCGGAACATGCCTCATGTGGC

GGCCAGGACGGCCAGCCGGGATCGGGATACTGGTCGTTACCAGAGCCACCGACCCGAGCA

AACCCTTCTCTATCAGATCGTTGACGAGTATTACCCGGCATTCGCTGCGCTTATGGCAGA

GCAGGGAAAGGAATTGCCGGGCTATGTGCAACGGGAATTTGAAGAATTTCTCCAATGCGG

GCGGCTGGAGCATGGCTTTCTACGGGTTCGCTGCGAGTCTTGCCACGCCGAGCACCTGGT

CGCTTTCAGCTGTAAGCGTCGCGGTTTCTGCCCGAGCTGTGGGGCGCGGCGGATGGCCGA

AAGTGCCGCCTTGCTGGTTGATGAAGTACTGCCTGAACAACCCATGCGTCAGTGGGTGTT

GAGCTTCCCGTTTCAGCTGCGTTTCCTGTTTGCCAGCCGGCCCGAGATCATGGGGTGGGT

GCTGGGCATCGTTTACCGCGTCATTGCCACGCACCTGGTCAAGAAAGCGGGCCATACCCA

CCAAGTGGCCAAGACGGGCGCGGTCACCCTGATCCAGCGTTTTGGATCGGCGCTCAATCT

GAATGTTCACTTCCACATGCTGTTTCTCGACGGTGTGTATGTCGAGCAATCCCACGGCTC

AGCGCGTTTCCGCTGGGTCAAGGCGCCGACCAGCCCAGAGCTCACCCAGCTGACGCACAC

CATCGCCCACCGGGTGGGTCGCTATCTGGAACGGCAAGGCCTGCTGGAACGGGATGTCGA

AAACAGCTATCTGGCCTCGGATGCGGTGGATGACGACCCGATGACACCCCTGCTGGGGCA

CTCGATCACTTACCGTATCGCTGTCGGTTCACAGGCGGGGCGAAAGGTGTTCACTTTGCA

AACTCTGCCGACCAGTGGTGATCCGTTCGGTGACGGGATTGGCAAGGTAGCCGGGTCCAG

CCTGCACGCCGGCGTGGCGGCCAGGGCCGATGAACGCAAGAAGCTCGAACGGCTGTGCCG

GTACATCAGCCGCCCGGCGGTATCCGAGAAGCGGCTGTCGTTAACACGAGGCGGCAACGT

GCGCTACCAGCTCAAGACGCCGTACCGGGACGGCACCACGCACGTCATTTTCGAACCATT

GGATTTCATTGCAAGGCTGGCCGCCCTGGTACCGAAGCCCAGAGTCAACCTAACCCGCTT

CCACGGGGTGTTCGCACCCAACAGTCGGCACCGGGCGTTGGTCACGCCGGCAAAACGGGG

CAGGGGCAACAAGGTCAGGGTGGCTGATGAACCGGCAACACCAGCACAACGGCGAGCGTC

GATGACATGGGCGCAACGGCTCAAGCGTGTTTTCAATATCGACATCGAGACCTGCAGCGG

CTGCGGCGGCGCCATGAAAGTCATCGCCTGCATTGAAGACCCTATAGTGATCAAGCAGAT

CCTTGATCACCTGAAGCACAAAGCCGAAACCAGCGGGACCAGGGCGTTACCCGAAAGCCG

GGCGCCACCGGCTGAGCTGCTCCTGGGTCTGTTTGACTGACGAGCCTGAAGGCCAACGAT

ACCAATCAAAATGCTGCGTTCACAGCGCCGCGGCAGGGATCCGCCGTGCTGGTTGTCGGA

AAAGGAGCCGCTAGTGGGAAAGAGGAGGGTAAATTTTCAGCGTTGCTGGCTCCCCGTCAG

CCGGATTGGGTTGCATCGCAGGGGTGTCGAAAGAGTCAACTGCGGTCCAAAGCTGTTGGA

CTTGGGTGAAAAGGGCGTTTATTCTTCCTATACGTTGTCGGCAGCGGGCCAAAAAGGAAT

ACGTCCATGCCCATCGAGGTGAAACCGGCTGTGAGCGCGGGTTCAAGCATATAGCCCGAC

AGGCGCGTATCCTTGCCGATCACGACACGATGGCGGTGGTCACCGCGACGAAAGACACGG

CCAGCCGCCATGCCGACGCGCAAGGCGGTTTCC

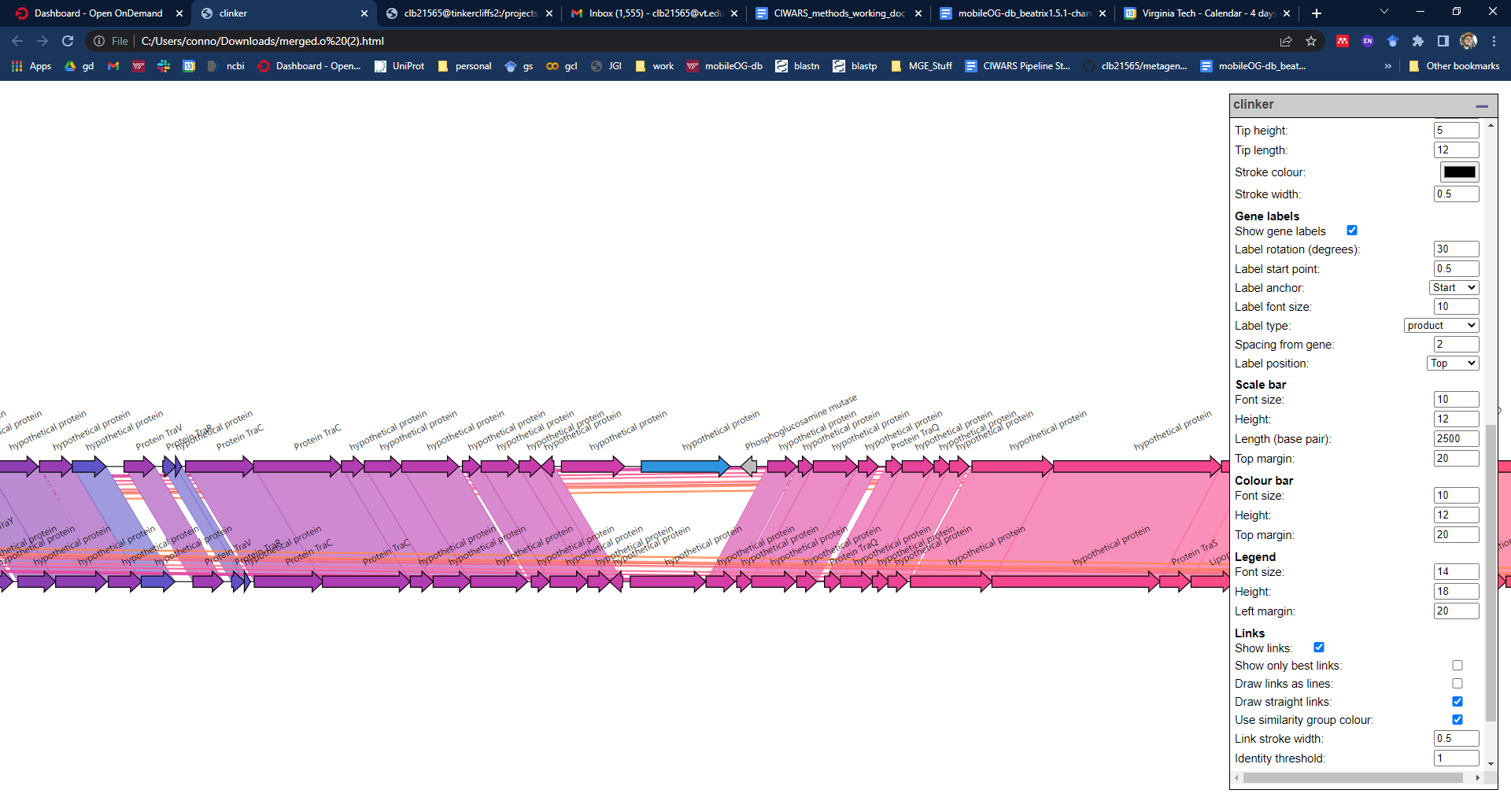

#### Fig. S2. A copy of an IS91 insertion sequence was extracted from a contig and used to create *in silico* insertions in the plasmid sequences for evaluating Kairos assess. (Left): nucleotide sequence of the associated insertion sequence predicted by ISEscan. (Right): Example insertion of IS91 copy into a plasmid assembly visualized using clinker. The top region is with the insertion and the bottom is the original plasmid sequence.

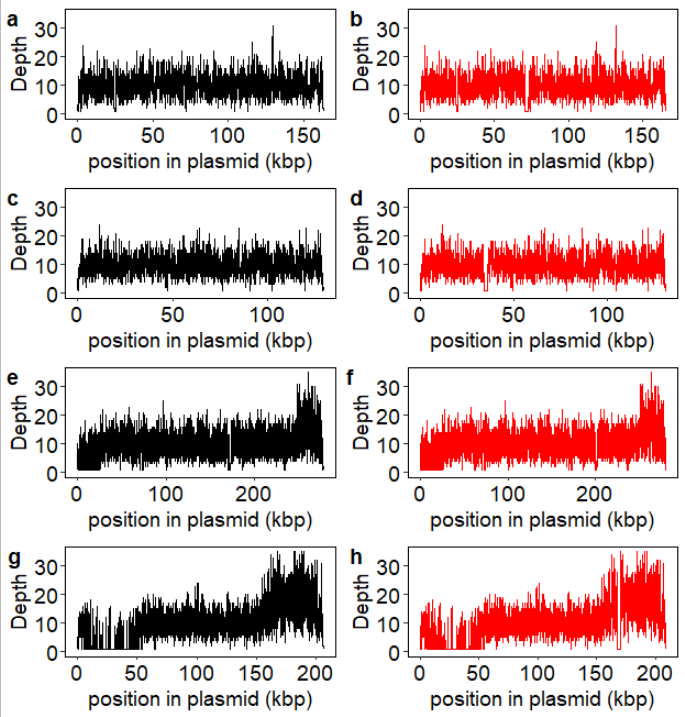

#### Figure S3. Depth profiles of *in silico* spike plasmid sequences. (a) *E. coli* plasmid without insert mapped with reads lacking insert sequences (black) (b) *E. coli* plasmid with insert mapped with reads lacking insert sequence (red).

#### Table S2. Spike-in experimental design for *in silico* evaluation of Kairos assess using the 10x coverage condition as an example.

| Table S2. Spike-in experimental design. | | | | | | | |
| --- | --- | --- | --- | --- | --- | --- | --- |
| Metagenomic Sample | HFGVI0044Q | UPKQF5024K | HDRFF4542I | JUJHF9134P | LNBYT5954Z | LZGQG4924W | EBZCS4093H |
| Spike-in | Ec_real_nospike_10x.fq.gz | Ec_real_nospike_10x.fq.gz | EC_plsdb_nospike_10x.fq.gz | EC_plsdb_nospike_10x.fq.gz | Ec_real_nospike_10x.fq.gz | Ec_real_nospike_10x.fq.gz | Ec_real_nospike_10x.fq.gz |
|  | Ar_real_nospike_10x.fq.gz | Ar_real_nospike_10x.fq.gz | Ar_plsdb_nospike_10x.fq.gz | Ar_plsdb_nospike_10x.fq.gz | Ar_real_nospike_10x.fq.gz | Ar_real_nospike_10x.fq.gz | Ar_real_nospike_10x.fq.gz |
|  |  | Ec_real_spike_10x.fq.gz |  | EC_plsdb_spike_10x.fq.gz | EC_plsdb_nospike_10x.fq.gz | EC_plsdb_nospike_10x.fq.gz | EC_plsdb_nospike_10x.fq.gz |
|  |  | Ar_real_spike_10x.fq.gz |  | Ar_plsdb_spike_10x.fq.gz | Ar_plsdb_nospike_10x.fq.gz | Ar_plsdb_nospike_10x.fq.gz | Ar_plsdb_nospike_10x.fq.gz |
|  |  |  |  |  | Ec_real_spike_10x.fq.gz | Ec_real_spike_10x.fq.gz | Ec_real_spike_10x.fq.gz |
|  |  |  |  |  | Ar_real_spike_10x.fq.gz | Ar_real_spike_10x.fq.gz | Ar_real_spike_10x.fq.gz |
|  |  |  |  |  |  | EC_plsdb_spike_10x.fq.gz | EC_plsdb_spike_10x.fq.gz |
|  |  |  |  |  |  | Ar_plsdb_spike_10x.fq.gz | Ar_plsdb_spike_10x.fq.gz |

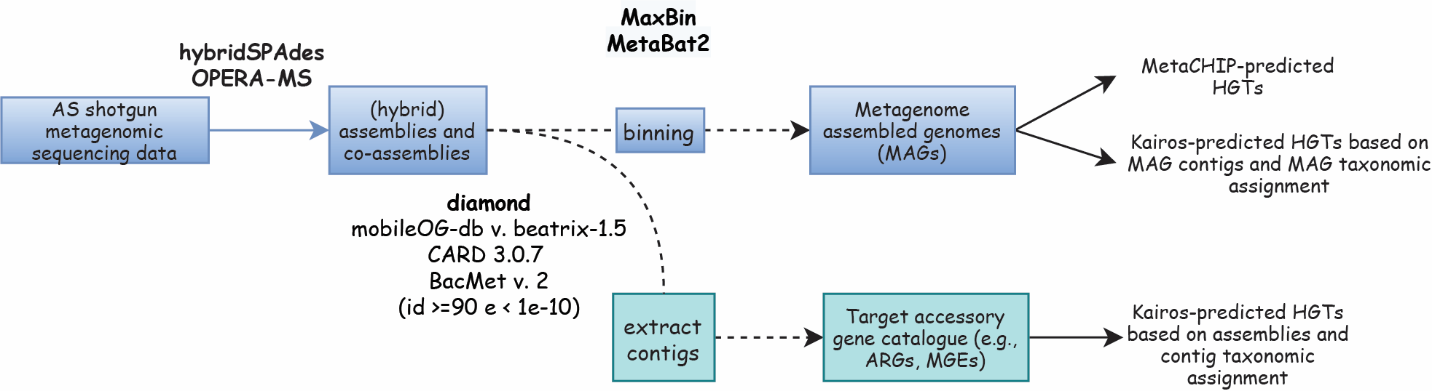

### Fig. S4. Experimental design comparing Kairos to MetaCHIP using data from Brown et al. 2023.

### Table S3. ARGs in Bin86, *Escherichia coli* D.

| Table S3. ARGs in Bin86, *Escherichia coli* D. | | | | | |
| --- | --- | --- | --- | --- | --- |
| bin | orf | stitle | pident | bitscore | evalue |
| bin86 | 23 | AAG57600.1\|FEATURES\|mphB\|macrolide_antibiotic\|mphB | 94.7 | 178.7 | 1.20E-46 |
| bin86 | 405 | AAC74000.1\|FEATURES\|msbA\|nitroimidazole_antibiotic\|msbA | 99.8 | 1063.5 | 3.40000000000071e-312 |
| bin86 | 571 | ABV18113.1\|FEATURES\|mdtG\|fosfomycin\|mdtG | 99.8 | 771.5 | 1.90E-224 |
| bin86 | 654 | NP_418574.1\|FEATURES\|Escherichia_coli_ampC_beta-lactamase\|cephalosporin\|Escherichia_coli_ampC_beta-lactamase | 98.4 | 768.5 | 1.50E-223 |
| bin86 | 669 | BAJ42218.1\|FEATURES\|Escherichia_coli_ampH_beta-lactamase\|cephalosporin\|Escherichia_coli_ampH_beta-lactamase | 99.2 | 766.5 | 5.60E-223 |
| bin86 | 686 | BAA15934.1\|FEATURES\|baeS\|aminoglycoside_antibiotic\|baeS | 99.6 | 913.7 | 3.50E-267 |
| bin86 | 687 | BAA15935.1\|FEATURES\|baeR\|aminoglycoside_antibiotic\|baeR | 100 | 484.2 | 3.50E-138 |
| bin86 | 698 | BAB35162.1\|FEATURES\|H-NS\|macrolide_antibiotic\|H-NS | 100 | 222.2 | 1.40E-59 |
| bin86 | 726 | BAE77933.1\|FEATURES\|CRP\|macrolide_antibiotic\|CRP | 99.5 | 418.7 | 1.60E-118 |
| bin86 | 830 | ACN32294.1\|FEATURES\|TolC\|macrolide_antibiotic\|TolC | 99.8 | 880.2 | 4.50E-257 |
| bin86 | 948 | BAE77781.1\|FEATURES\|mdtE\|macrolide_antibiotic\|mdtE | 99.5 | 720.3 | 4.60E-209 |
| bin86 | 1004 | AFH35853.1\|FEATURES\|Escherichia_coli_mdfA\|tetracycline_antibiotic\|Escherichia_coli_mdfA | 94.6 | 740 | 6.00E-215 |
| bin86 | 1387 | AAC76093.1\|FEATURES\|bacA\|peptide_antibiotic\|bacA | 100 | 468.8 | 1.50E-133 |
| bin86 | 1403 | AAC76298.1\|FEATURES\|AcrF\|fluoroquinolone_antibiotic\|AcrF | 99.5 | 1937.2 | 0 |
| bin86 | 1448 | BAA11237.1\|FEATURES\|emrY\|tetracycline_antibiotic\|emrY | 100 | 431.8 | 1.90E-122 |
| bin86 | 1542 | BAE78084.1\|FEATURES\|mdtN\|nucleoside_antibiotic\|mdtN | 100 | 641.3 | 2.40E-185 |
| bin86 | 1543 | BAE78083.1\|FEATURES\|mdtO\|nucleoside_antibiotic\|mdtO | 98.2 | 1315.4 | 0 |
| bin86 | 1553 | AAC74149.2\|FEATURES\|mdtH\|fluoroquinolone_antibiotic\|mdtH | 100 | 783.9 | 3.60E-228 |
| bin86 | 1878 | AAC75733.1\|FEATURES\|emrB\|fluoroquinolone_antibiotic\|emrB | 100 | 985.3 | 1.00E-288 |
| bin86 | 1879 | BAA16547.1\|FEATURES\|emrA\|fluoroquinolone_antibiotic\|emrA | 99 | 180.6 | 3.30E-47 |
| bin86 | 2059 | AAC73565.1\|FEATURES\|Escherichia_coli_acrA\|fluoroquinolone_antibiotic\|Escherichia_coli_acrA | 100 | 511.1 | 3.00E-146 |
| bin86 | 2197 | AAC75429.1\|FEATURES\|evgS\|macrolide_antibiotic\|evgS | 97.8 | 1917.9 | 0 |
| bin86 | 2308 | AAC75731.1\|FEATURES\|emrR\|fluoroquinolone_antibiotic\|emrR | 100 | 347.8 | 2.90E-97 |
| bin86 | 2388 | CBJ02047.1\|FEATURES\|Escherichia_coli_ampC1_beta-lactamase\|cephalosporin\|Escherichia_coli_ampC1_beta-lactamase | 99.1 | 883.6 | 3.60E-258 |
| bin86 | 2935 | BAE78116.1\|FEATURES\|eptA\|peptide_antibiotic\|eptA | 99.5 | 443 | 7.90E-126 |
| bin86 | 2966 | AAC75314.1\|FEATURES\|PmrF\|peptide_antibiotic\|PmrF | 99.7 | 628.6 | 1.50E-181 |
| bin86 | 3156 | AAC75137.1\|FEATURES\|mdtC\|aminocoumarin_antibiotic\|mdtC | 99.7 | 602.8 | 9.00E-174 |
| bin86 | 3205 | BAA16344.1\|FEATURES\|acrD\|aminoglycoside_antibiotic\|acrD | 98.8 | 460.3 | 5.40E-131 |

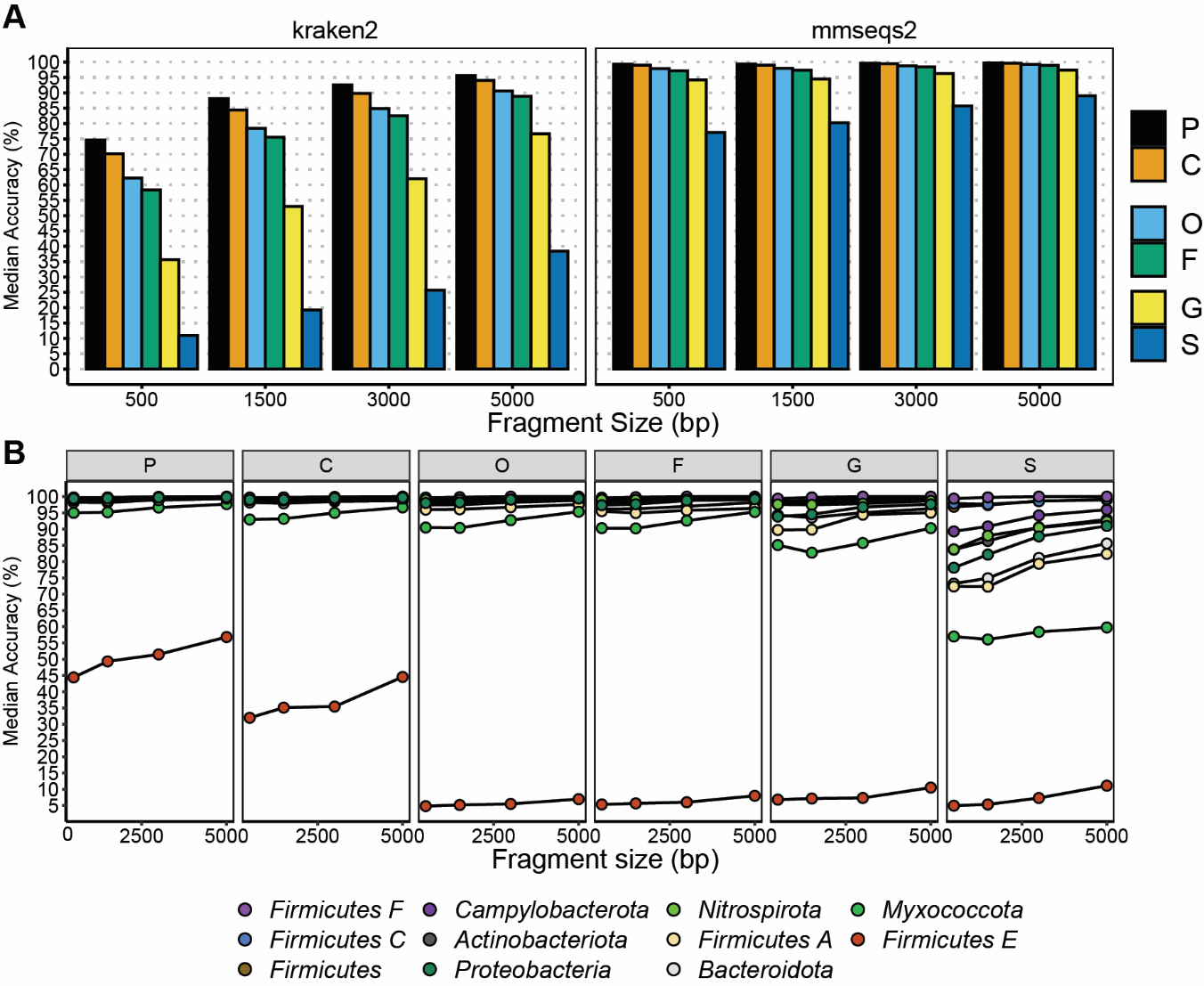

### Fig. S5. Contig taxonomy predictions faithfully capture many important environmental taxa. (A) Comparison of median accuracies across kraken2 and mmseqs2 taxonomy (both using gtdb). (B) mmseqs2 taxonomy predictions showed strong performance for may important environmental taxa that improved with increasing fragment length at higher taxonomic levels. *p:phylum; c:class; o:order; f:family; g:genus; s: species.*

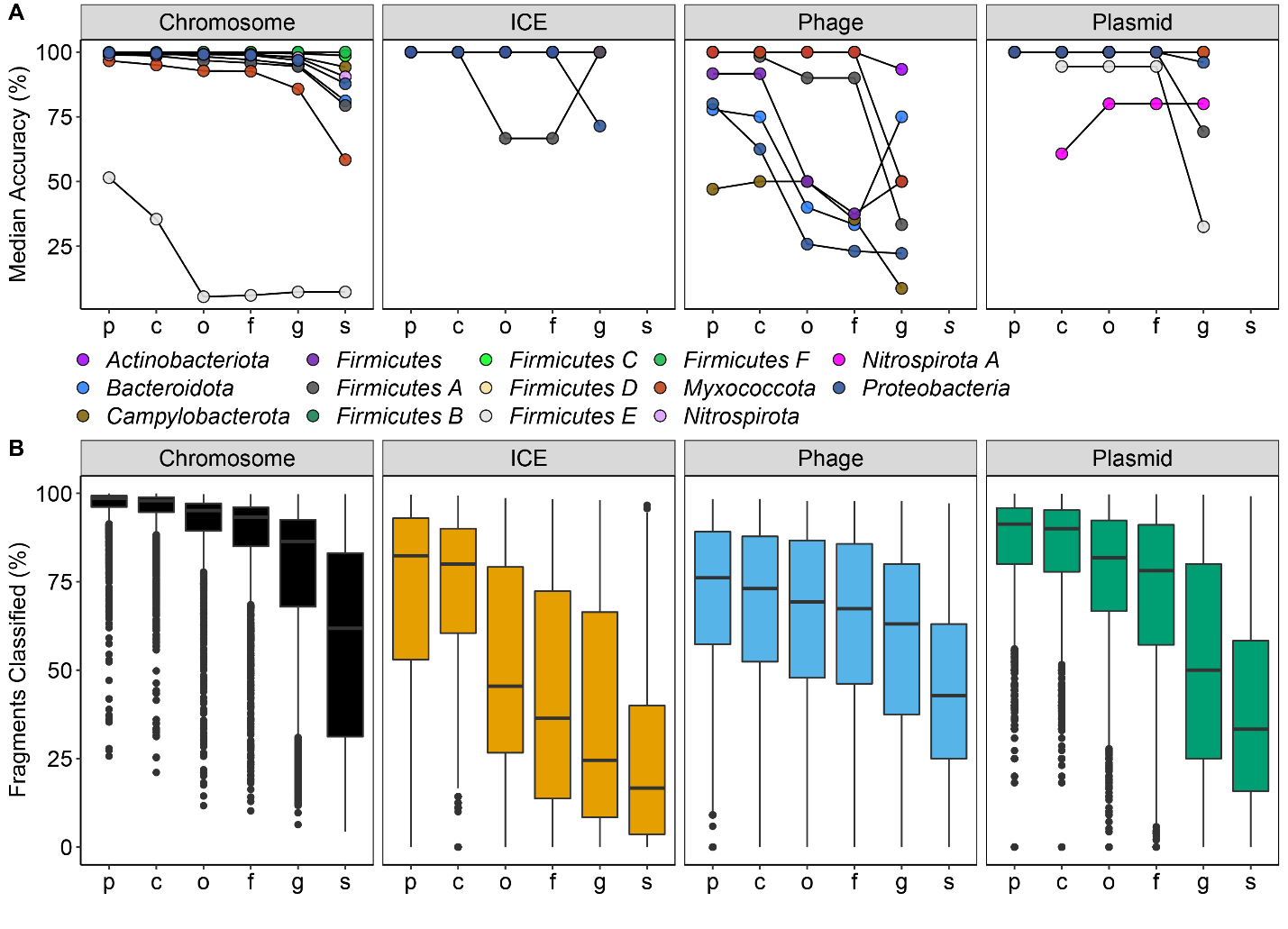

### Fig. S6. Taxonomic assignments by mmseqs2 faithfully reflect phyla of origin for chromosomes, integrative elements, and plasmids. (A) Select environmental-associated phyla and their median classification accuracies (considering all classified fragments). (B) Classification rates for different fragment types. *p:phylum; c:class; o:order; f:family; g:genus; s: species.*

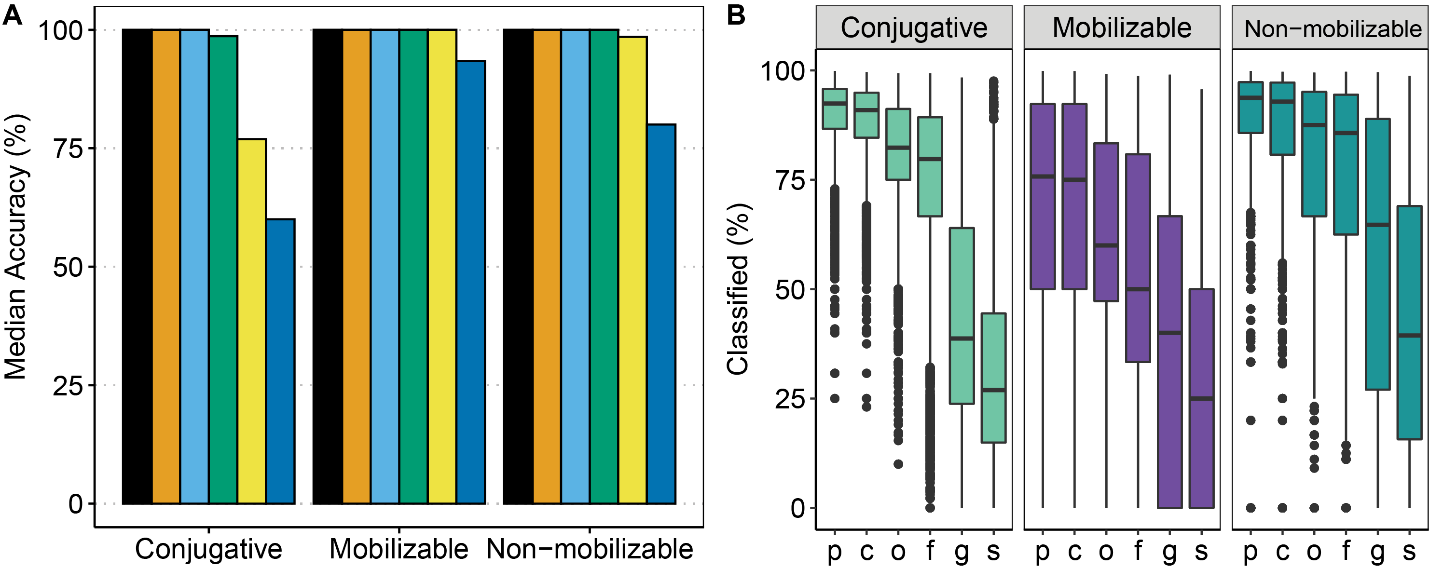

### Fig. S7. *Proteobacteria-*associated plasmids display disparate classification accuracies. (A) Median accuracy for classifications remained similar to aggregate values for MGEs and chromosomes although with worse performance for higher levels. (B) *Proteobacteria-*associated plasmids displayed poor classification rates. This was most substantial for conjugative and mobilizable plasmids. *p:phylum; c:class; o:order; f:family; g:genus; s: species.*

### Table S5. Result of *in silico* spike-in experiment.

| **Table S5. Result of *in silico* spike-in experiment.** | | | | | | | | |
| --- | --- | --- | --- | --- | --- | --- | --- | --- |
| exp3.1 (88% coverage criteria for mmseqs edge clustering) | | | | | | | | |
| 90% | Prediction | Reality | n |  | prp | prn |  |  |
| 1 | present | present | 34 | ap | 34 | 1 | precision | 1 |
| 2 | absent | absent | 21 | an | 0 | 21 | recall | 0.97 |
| 3 | present | absent | 1 |  |  |  |  |  |
| 50% | Prediction | Reality | n |  | prp | prn |  |  |
| 1 | present | present | 34 | ap | 34 | 0 | precision | 0.85 |
| 2 | absent | absent | 16 | an | 6 | 16 | recall | 1.00 |
| 3 | present | absent | 6 |  |  |  |  |  |
| exp3.0 (100% coverage criteria for mmseqs edge clustering) | | | | | | | | |
| 90% | Prediction | Reality | n |  | prp | prn |  |  |
| 1 | absent | absent | 22 | ap | 22 | 22 | precision | 1.00 |
| 2 | present | present | 22 | an | 0 | 12 | recall | 0.50 |
| 3 | absent | present | 12 |  |  |  |  |  |
| 80% | Prediction | Reality | n |  | prp | prn |  |  |
| 1 | present | present | 33 | ap | 33 | 19 | precision | 0.92 |
| 2 | absent | absent | 19 | an | 3 | 1 | recall | 0.63 |
| 3 | present | absent | 3 |  |  |  |  |  |
| 4 | absent | present | 1 |  |  |  |  |  |
| 50% | Prediction | Reality | n |  | prp | prn |  |  |
| 1 | present | present | 34 | ap | 34 | 0 | precision | 0.89 |
| 2 | absent | absent | 18 | an | 4 | 18 | recall | 1.00 |
| 3 | present | absent | 4 |  |  |  |  |  |

*Percentages refer to the proportion of unique loci of a given variant detected. *prp:*predicted positive; *prn:*predicted negative; *ap:* actual positive; *an:* actual negative.
